# Natural selection and evolution of mitochondrial genome in terrestrial and aquatic turbellarians

**DOI:** 10.1101/2024.05.12.593061

**Authors:** Rong Li, Hui Feng, Pinglin Cao, Jianxing Wang

## Abstract

Mitochondrial energy metabolism may be directly influenced by natural selection pressures in response to the energy demands of an organism’s specific ecological niche. Here, we hypothesized that the mitochondrial genome of turbellarian animals was influenced by the metabolic requirements of various habitats We used selection pressure and phylogenetic independent contrasts (PIC) analyses to detect the selection pressure of protein-coding genes in the mitochondrial genome of turbellarian living in aquatic and terrestrial habitats. The results indicated turbellarians in aquatic habitats experienced greater selection pressure, resulting in the evolution of several genes at a lower ω value compared to their terrestrial counterparts. The NAD4 was identified as having positive selection sites in all site models analyzed. Among all genes, the equilibrium constant (ionization of COOH) was the most frequently detected amino acid characteristic with significant positive selection changes, followed by hydropathy and molecular volume. The equilibrium constant (ionization of COOH) was found to be the amino acid characteristic with the greatest magnitude of change in NAD4. This suggests that it plays a crucial role in the adaptive evolution of turbellarians to varying habitats. The study found a significant positive correlation (R = 0.61, p < 0.05) between the ω value of turbellarian and their habitats by PIC analysis. These findings shed light on the adaptive evolution of turbellarian mtDNAs and their influence on the oxidative phosphorylation molecular mechanism.

**Significance statement:** The adaptive evolution of mitochondrial genes is closely related to environmental temperature, high energy metabolism demand, altitude, and oxygen availability. Mitochondrial energy metabolism may be directly influenced by natural selection pressures in response to the energy demands of an organism’s specific ecological niche. Here, we investigated the evolution of mtDNA PCGs in turbellarians across different habitats, and identified purifying selection as the main evolutionary pattern of the turbellarian mitochondrial genome. Additionally, we found that NAD4 played an important role in the adaptive evolution of turbellarians to different habitats.

## 1. Introduction

Mitochondria encode 37 genes capable of producing roughly 95% of ATP through the processes of oxidative phosphorylation or electron transfer[1]. Thirteen protein-coding genes (PCGs) of these encode core structures of four protein complexes (complexes I, III, IV, and V) that directly participate in electron transfer and ATP synthesis. These two processes are thought to have played a significant role in the evolution of the mitochondrial genome[2]. Moreover, mitochondrial oxidative phosphorylation produces both energy and heat, which must be balanced with ATP synthesis. The temperature of the environment may limit the genetic level ability of oxidative phosphorylation[3, 4]. Subsequently, genes in the mitochondrial genome which involved in electron transfer, may play a crucial adaptive role in the evolution of organisms[5]. Investigations have found that the adaptive patterns of organisms in response to ecological changes are more diverse than expected[6], and thus, the adaptive evolution of mitochondrial genomes to different environmental conditions has become a research hotspot in evolutionary biology. The selective pressure may directly act on mitochondrial energy metabolism to drive the adaptation of organisms to the different energy demands required by their unique ecological niche[7]. Extreme environments may promote adaptive evolution of different mitochondrial genes in certain lineages[8]. The adaptive evolution of mitochondrial genes is closely related to environmental temperature, high energy metabolism demand, altitude, and oxygen availability[5, 9]. Especially in extreme environmental conditions, the molecular evolution of mitochondrial genomes in high-altitude regions has attracted much attention[8, 10]. This is because by studying the selection pressure of environmental temperature and oxygen availability on the molecular changes of the mitochondrial genome, we can gain important insights into the adaptive evolution of mitochondrial genomes[10].

In nature, mutations in genes often have no effect on the organisms or only produce minor detrimental effects. Only when a mutation is beneficial to organisms can it be preserved through positive selection. Extreme environmental changes can have a significant impact on the metabolism of organisms, as changes in metabolic capacity are typically directly related to mutations in genes encoding proteins involved in oxidative phosphorylation. For instance, the NADH dehydrogenase subunit gene can produce a proton pump[11],while the ATP synthase subunit gene can affect ATP synthesis [12].Similarly, the cytochrome b gene can catalyze the proton transfer coupled from ubiquinone to cytochrome c[13], and the cytochrome C oxidase subunit gene directly participates in electron and proton transfer [14]. Mutations in these protein-coding gene loci can affect the metabolic capacity of organisms. It has been demonstrated that the mitochondrial genome undergoes adaptive evolution, and studies on the mitochondrial genome of small brown planthopper suggest that beneficial gene mutations may increase their cold tolerance and reproductive ability, and bring genetic advantages[15].

The turbellarians were belonging to the phylum Platyhelminthes and the class Rhabditophora, which includes all incompletely parasitic subgroups of Platyhelminthes, comprising approximately 4,500 species[16]. Most of them live in the ocean and crawling on the seabed mud, rocks, and algae, while some are planktonic or symbiotic on the external surfaces of marine animals, some species are distributed in freshwater environments or on humid land in tropical and subtropical areas[17]. As the energy demand for movement is the primary determinant of metabolic rate[18], the different lifestyles of turbellarians are expected to affect their energy requirements, resulting in different metabolic rates. Therefore, we tested the mtDNA selection pressure among turbellarian animals living in different habitats using the rate ratio of non-synonymous/synonymous nucleotide substitutions (dN/dS (ω)) and determine whether there is an adaptive molecular basis for mitochondrial selection pressure among turbellarian animals living in different environments.

## 2. Materials and Methods

### 2.1. Data Acquisition

All data used in this study were obtained from the National Center for Biotechnology Information (NCBI) database. The complete mitochondrial genomes of 25 species of turbellarian were retrieved and downloaded from the NCBI database. Of these, 14 species were found in aquatic habitats while 11 species were found in terrestrial habitats. The accession numbers and mitochondrial genome sizes for all species included in the study are recorded in Table S1.

### 2.2. Phylogenetic reconstruction

In this study, the sequences of 12 PCGs from the mitochondrial genomes of all species (the ATP8 was missing in most of the studied species) were extracted using PhyloSuite v1.2.2 [19] software, and the nucleotide sequences of each PCG were individually aligned using MAFFT v7.313[20], after poorly aligned positions were eliminated using Gblocks v0.91b [21] with default parameters. The individually aligned sequences were then concatenated to generate a combined sequence dataset. We reconstructed a maximum likelihood phylogenetic tree using IQ-TREE software[22], and performed 1000 ultrafast bootstrap replicates to evaluate node support, with only nodes having bootstrap values > 95 considered reliable. The resulting phylogenetic tree was visualized in iTOL [23].

### 2.3. Selection pressure analyses

The level of natural selection pressure on PCG can be measured by the value of dN/dS (ω), where dS represents the synonymous rate and dN represents the non-synonymous rate. When there is no selection pressure, the synonymous replacement rate and the non-synonymous replacement rates are equal, at which point dN/dS = 1. When subject to negative or purification selection pressure, natural selection will prevent amino acids from changing, and the synonymous replacement rate will be greater than the non-synonymous replacement rate, i.e., dN/dS < 1; Conversely, under positive selection pressure, the rate of amino acid substitution is favored by natural selection, leading to potential adaptive changes in protein function, resulting in a dN/dS ratio greater than 1.

### 2.4. Branch model

To test whether the ω values differ between aquatic and terrestrial turbellarians, we used the branch model of CodeML program in the PAML [24] to analyze the ω values of 12 PCGs from 24 species. This model is mainly used to detect whether the ω value on a specific branch is significantly higher than the background branch, that is, whether the gene evolves faster on the target branch. The one-ratio model (model = 0, NSsites = 0) was first used to estimate a unique ω value for each PCG on all branches of the phylogenetic tree. Then, assuming that each branch has an independent ω value, the free-ratio model (model = 1, NSsites = 0) was used to estimate the ω values of each PCG in 24 species, to evaluate the selection pressure on the terminal branches. Finally, the two-ratio model (model = 2, NSsites = 0) was used to assume that aquatic and terrestrial turbellarians have different ω values, and aquatic turbellarians were marked as the foreground branch of the phylogenetic tree for the two-ratio model and the all branch site model described below. The remaining terrestrial turbellarians that were not marked were considered as background branches by the program. The Hyphy software [25] RELAX program [26] was used with the same branch label scheme to infer whether the selection strength in aquatic turbellarians was enhanced (K > 1) or relaxed (K < 1). If the dN or dS value equals 0, the ω value can be smaller or larger, and this study did not use these ω values in the analysis, but assigned them as NA data in the ω dataset. The Wilcoxon test was used to evaluate the statistical significance of the differences in ω values of 12 PCGs among different habitat species.

### 2.5. Branch site model

The branching site model was mainly used to detect the presence of positive selection site for genes on foreground branches, and the model assumes that the ω values were variable between sites and also between branches. The aquatic turbellarians were used as foreground branches and terrestrial turbellarians as background branches, and the likelihood ratio test (LRT) was used to compare the null hypothesis (Model A null) with the alternative hypothesis (Model A). The posterior probability of positive selection site was calculated by Bayes Empirical Bayes (BEB) method, and the presence of a probability value > 0.95 positive selection site indicated that the gene was under positive selection pressure on the target branch. Finally, another branch site analysis was applied in this study using aBSREL (adaptive branch-site random effects likelihood) from HyPhy [25].

### 2.6. Site model

The site model was primarily used to detect positive selection site in genes. During the analysis, the ω value of each branch in the evolutionary tree was assumed to be the same, and two models were compared: The first model was the null model, assuming that all sites have ω values <1 or = 1. The second was the positive selection model, allowing for ω values <1, = 1, or > 1 at certain sites. The likelihood ratio test (LRT) was used to compare the difference in the likelihood values (lnL) between the two models, and a p value was calculated using a chi-square test (with 2 degrees of freedom). If p < 0.05, the null model is rejected and positive selection sites are inferred [24]. In addition, it is recommended to use Model 7 and Model 8 for comparison, which divide the ω values into 10 categories and have more relaxed p value results than other comparison methods, allowing for the detection of more positively selected genes. The PAML site model can detect positive selection sites in genes at the overall level, but cannot indicate whether a gene is under positive selection pressure on a specific evolutionary branch [24].

In addition to the site model analysis with CodeML, we also used the following programs in the Hyphy software [25] to infer positively selected codons: FUBAR (Fast, Unconstrained Bayesian AppRoximation) [27], MEME (Mixed Effects Model of Evolution) [28], and FEL (Fixed Effects Likelihood) [29]. For each method, we applied the best fitting substitution model to each gene and evaluated significance based on a posterior probability > 0.9 (FUBAR) or p < 0.05 (MEME) [2].

We also used the TreeSAAP software [30] to evaluate significant changes in amino acid features to explore their roles in the process of phylogenetic evolution. When amino acid substitutions significantly affect the biochemical properties of proteins, they were considered as candidate sites under selection. Analysis was performed using a sliding window of 20 codons and a step of 1 codon. Amino acid replacements were considered as possibly under selection when the degree of change was ≥ 6. At the same time, Z-scores greater than 3.09 or less than −3.09 (p <0.001) were attributed to positive and purifying selection, respectively [31].

### 2.7. Phylogenetic independent contrasts analysis

The close relationship between species tends to result in similarities, that is, shared inheritance, which is known as phylogenetic inertia and may affect inter-species comparative analysis [32]. To eliminate the influence of evolutionary inertia and explore the relationship between different habitats and the ω values of mitochondrial PCGs, we used Phylogenetic Independent Contrast (PIC) [33] analysis in the ape package in R to eliminate the influence of closely related species (i.e. shared inheritance). Firstly, the turbellarians were divided into aquatic and terrestrial turbellarians according to their different habitats, which were coded as 0 and 1, respectively. Then, the ω value of each species was obtained from the branch model analysis with CodeML, and the tree file used for the analysis was obtained from the Maximum Likelihood tree obtained from the above phylogenetic analysis.

## 3. Result

### 3.1. Phylogenetic reconstruction

The nucleotide sequences of 12 PCGs from mitochondrial genomes were used to reconstruct the phylogenetic relationships among turbellarians. The maximum likelihood method reconstructed well-supported phylogenetic relationships for the turbellarians with high bootstrap support for most branches (Fig. 1). The 11 terrestrial turbellarians all belonged to the family Geoplanidae and clustered together with a bootstrap value of 99.8, while the remaining 13 aquatic turbellarians were distributed in different taxonomic groups (Tricladida: 6; Rhabdocoela: 2; Polycladida: 4; Macrostomida: 1) on the phylogenetic tree.

**Figure 1.**
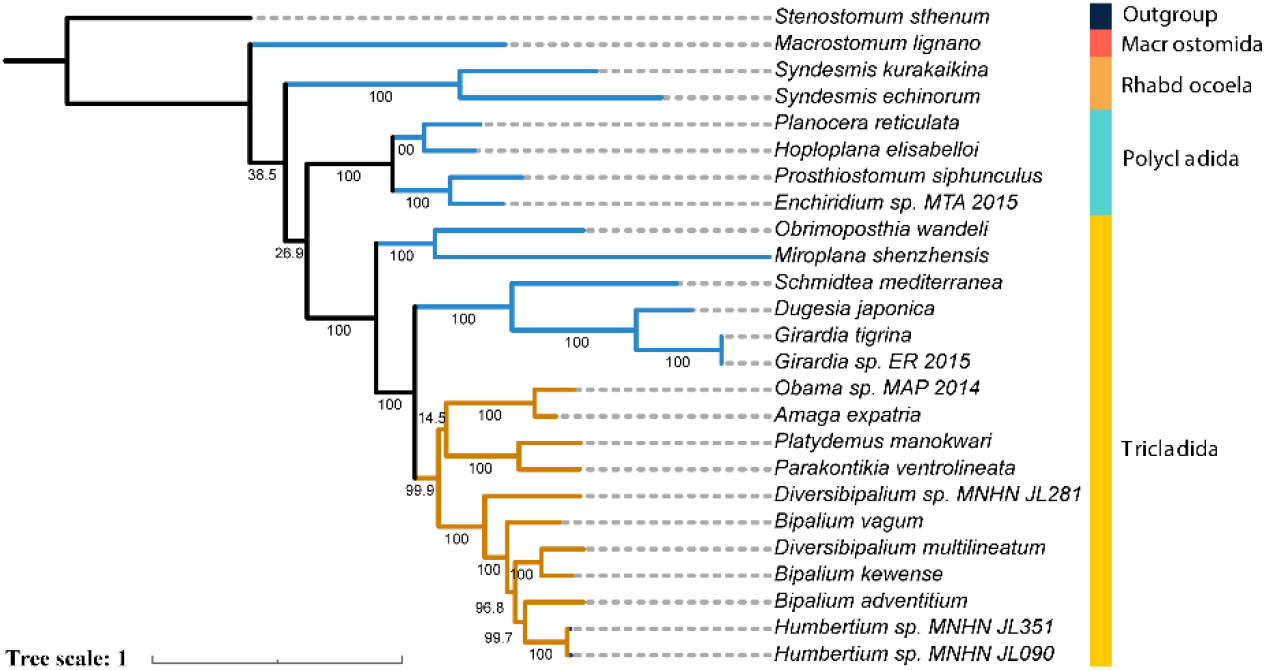
Phylogenetic tree based on 12 protein-coding genes (A: Maximum likelihood, B: Bayesian inference). Brown branches (Terrestrial species), Blue branches (aquatic species), black branches (outgroup).

### 3.2. Branch model

To test the role of natural selection in the 12 PCGs of the turbellarians, we used the branch model of CodeML. As shown in Table S2, for all 12 PCGs of the turbellarians, the free-ratio model was more suitable for the data than the one-ratio model, and it was found that the PCGs of the 24 turbellarians experienced different selection pressures in different lineages. When aquatic turbellarians were marked as foreground branches, the two-ratio model was used to calculate the selection pressure in different lineages, branch model analysis revealed that compared to aquatic turbellarians, three genes were identified experienced higher selection pressures in aquatic turbellarians, which were ATP6 (terrestrial: ω, 0.04155; aquatic: ω, 0.02375), NAD2 (terrestrial: ω, 0.06412; aquatic: ω, 0.02258), and NAD4L (terrestrial: ω, 0.07146; aquatic: ω, 0.01658), as well as a slowly evolving gene, COX1 (terrestrial: ω, 0.02463; aquatic: ω, 0.03408)(Fig. 2). In addition, the RELAX program of the Hyphy software can infer whether the selection strength in the PCGs of aquatic turbellarians has been enhanced (K > 1) or relaxed (K < 1). As shown in Table 1, compared to terrestrial turbellarians, significant intensified selection (K > 1) was detected in ATP6, NAD1, NAD3, NAD4, NAD4L, NAD5, and CYTB of aquatic turbellarians, with only COX1 under relaxed selection, but the result was not significant (K = 0.94, p = 0.524).

**Table 1.** The RELAX program tests for genes that have significantly enhanced selection.

| Gene | K | LR | p-value | Significant |
| --- | --- | --- | --- | --- |
| ATP6 | 1.66 | 33.53 | 0 | ** |
| COX1 | 0.94 | 0.41 | 0.524 | no |
| COX2 | 1.21 | 3.22 | 0.073 | no |
| COX3 | 1.17 | 3.33 | 0.068 | no |
| NAD1 | 1.26 | 12.23 | 0 | ** |
| NAD2 | 1.06 | 0.47 | 0.493 | no |
| NAD3 | 2.29 | 6.8 | 0.009 | ** |
| NAD4 | 1.15 | 11.52 | 0.001 | ** |
| NAD4L | 2.57 | 15.69 | 0 | ** |
| NAD5 | 1.68 | 34.29 | 0 | ** |
| NAD6 | 1.04 | 0.37 | 0.545 | no |
| CYTB | 1.29 | 10 | 0.002 | ** |

**Figure 2.**
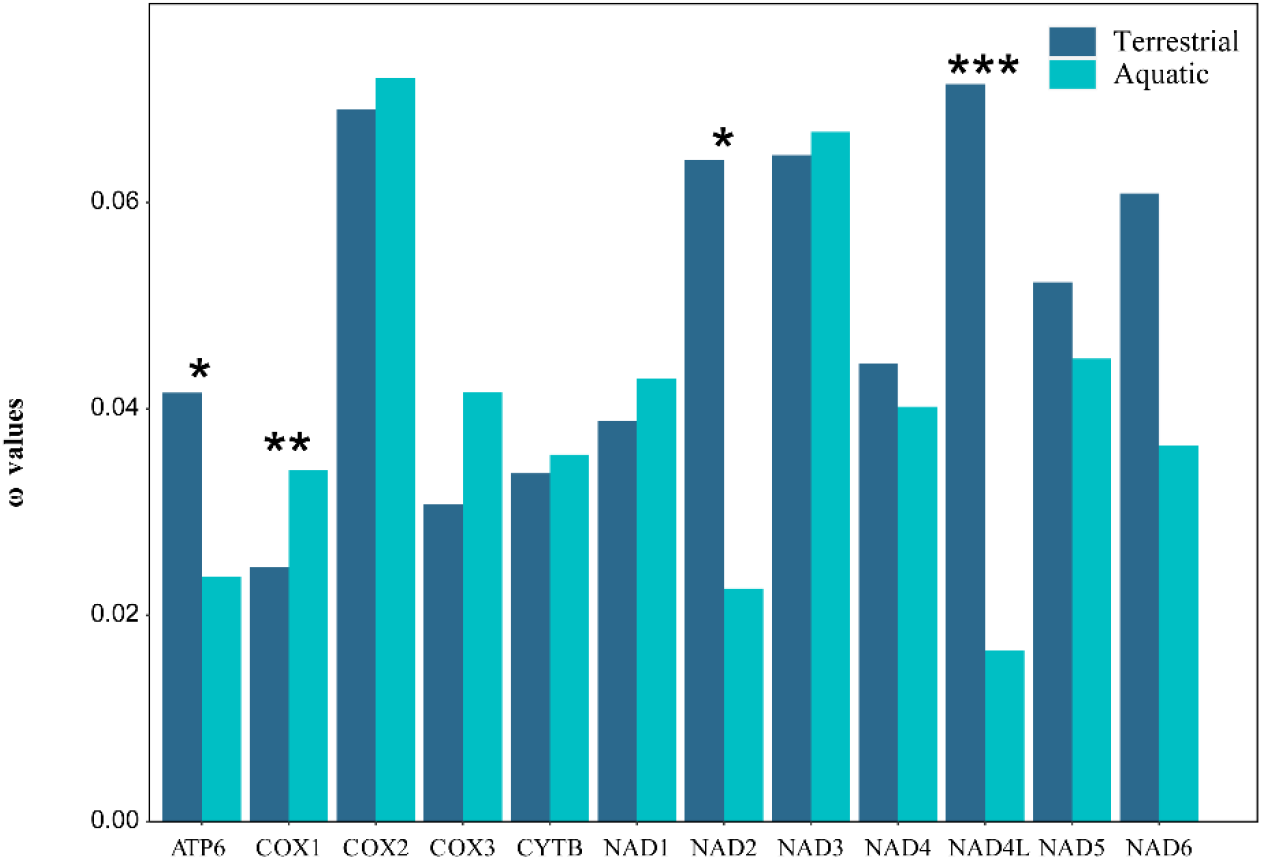
The ω values of 12 protein-coding genes in terrestrial and aquatic flatworms under two-ratio model (*\* p* < 0.05, *\*\* p* < 0.01, *\*\*\* p* < 0.001)

The free-ratio model assigns a separate ω value to each branch, when comparing ω values based on each PCG for each species, terrestrial turbellarians had higher ω values than aquatic turbellarians in ATP6, COX2, CYTB, NAD1-2, NAD4-6, and NAD4L, although this difference was significant only in NAD4 (p < 0.05) (Fig. 3).

**Figure 3.**
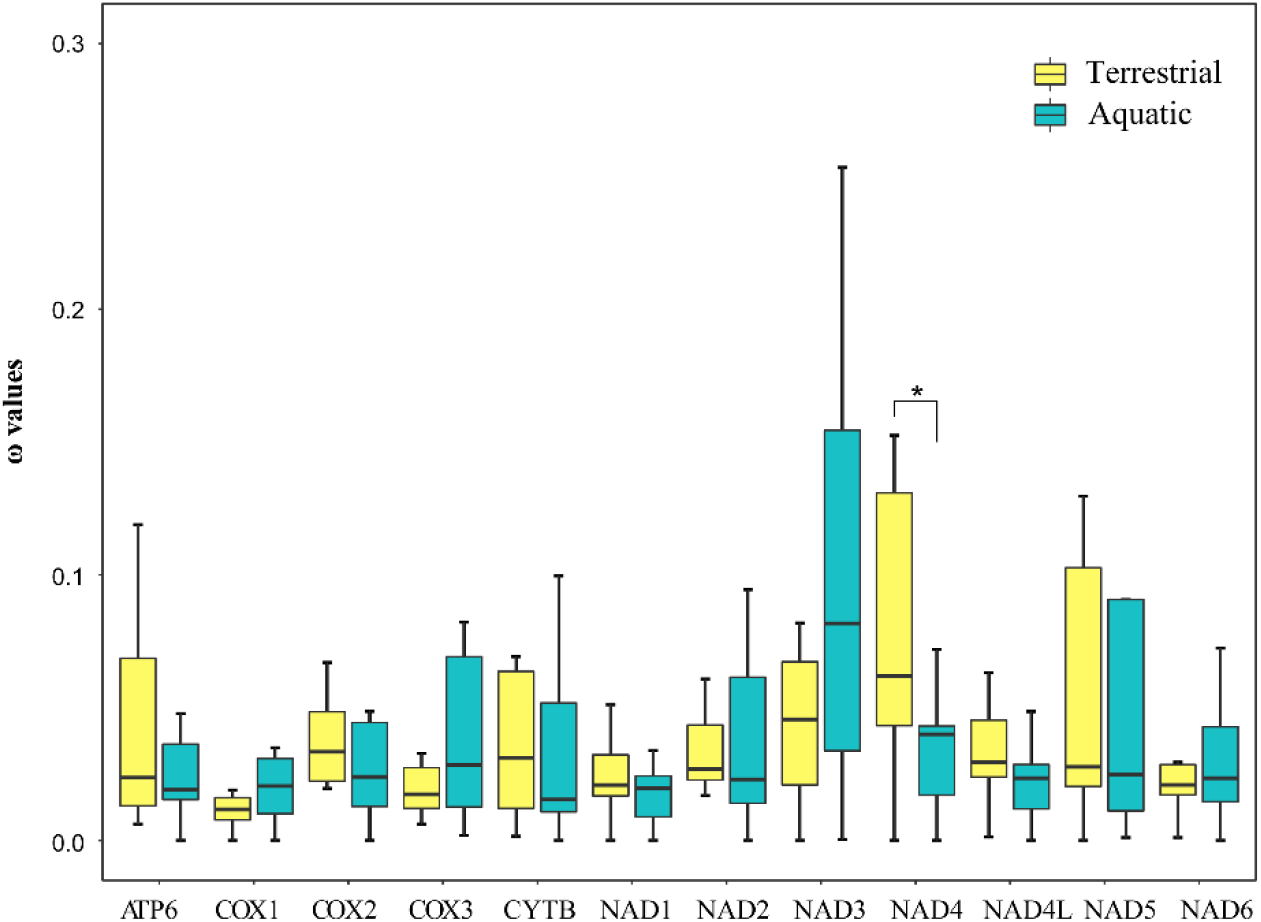
Comparison of ω values of each protein-coding gene between terrestrial and aquatic flatworms under the free-ratio model

### 3.3. Branch site model

We applied codeML to conduct branch-site tests on each PCG of aquatic and terrestrial turbellarians. Multiple significant signals of positive selection were detected in the PCGs of aquatic turbellarians, ATP6, COX1, COX2, and COX3 had the most positive selection sites, while no significant positive selection sites were detected in NAD4L and NAD6 (Table S3). Based on aBSREL analysis, three branches of turbellarians experienced episodic diversifying selection, 2.3% sites of NAD2 underwent positive selection in the lineage M. lignano, 2.2% sites of NAD4 underwent positive selection in S. kurakaikina, and 4.4% sites of CYTB underwent positive selection in D. japonica and M. lignano (Table 2).

**Table 2.** Codon positions under positive selection detected by branch-site model using aBSRE.

| Gene | % of positive selected sites | Lineage |
| --- | --- | --- |
| ATP6 | NA |  |
| COX1 | NA |  |
| COX2 | NA |  |
| COX3 | NA |  |
| NAD1 | NA |  |
| NAD2 | 2.30% | *Macrostomum lignano* |
| NAD3 | NA |  |
| NAD4 | 2.20% | *Syndesmis kurakaikina* |
| NAD4L | NA |  |
| NAD5 | NA |  |
| NAD6 | NA |  |
| CYTB | 4.40% | *Dugesia japonica, Macrostomum lignano* |

### 3.4. Site model

To detect positive selection sites in the PCGs of the turbellarians at the global level, we used the site model of CodeML and the significance results were compared the likelihood ratio test (LRT) between two models, Model 7 and Model 8, and test using chi-square test. Positive selection sites were detected in several genes, with position 469 Q in COX1, 81 N in CYTB, 77 F in NAD2, 77 R in NAD4, and 11 I and 14 F in NAD6 (Table 3). However, only the 77R position in NAD4 was validated by the chi-square test (p = 0.0043178) (Table 3).

**Table 3.** Codon positions under positive selection detected by site model using CodeML, MEME, FUBAR and FEL.

| Gene | CodeML | MEME | FUBAR | FEL |
| --- | --- | --- | --- | --- |
|  | Positive selected sites | Positive selected sites | Positive selected sites | Positive selected sites |
| ATP6 | NA | 72 *, 154 * | NA | NA |
| COX1 | 469 Q | 272 *, 526 * | NA | 608 * |
| COX2 | NA | 45 *, 300 *, 319 * | NA | NA |
| COX3 | NA | 15 *, 301 * | NA | 15 * |
| NAD1 | NA | NA | NA | NA |
| NAD2 | 77 F | 161 *, 223 * | NA | 161 *, 197 *, 210 *, 223 * |
| NAD3 | NA | 81 * | NA | NA |
| NAD4 | 77 R ** | 45 *, 316 * | 87 * | 316 * |
| NAD4<br>L | NA | NA | NA | NA |
| NAD5 | NA | 140 *, 320 * | NA | NA |
| NAD6 | 11 I, 14 F | NA | 175 * | NA |
| CYTB | 81 N | 39 *, 140 *, 235 * | NA | 421 * |

We also detected more significant positively selected codons based on the MEME, FUBAR, and FEL algorithms compared to CodeML. Specifically, 19 positively selected codons were detected in 9 genes (ATP6, COX1-3, CYTB, and NAD2-5) based on MEME, 2 positively selected codons were found in NAD4 and NAD6 using the FUBAR algorithm, and 8 positively selected codons were identified in 5 genes (COX1, COX3, CYTB, NAD2, and NAD4) based on FEL (Table 3). Among them, the site 15 in COX3, site 161 and 223 in NAD2, and site 316 in NAD4 were detected by two algorithms, while the remaining positively selected codons could only be detected by one of the three algorithms.

We also used the TreeSAAP software to investigate the degree of impact of amino acid substitutions on the physicochemical properties of proteins, and discover significant positive changes in the amino acid characteristics at 87 sites in the 12 PCGs of the turbellarians, and purifying selection was found to be widespread in all 12 genes (Table 4). The 87 sites included 8 sites in ATP6, 6 sites in COX1, 10 sites in COX2, 9 sites in COX3, 6 sites in CYTB, 8 sites in NAD1, 4 sites in NAD2, 5 sites in NAD3, 10 sites in NAD4, 2 sites in NAD4L, 11 sites in NAD5, and 8 sites in NAD6 (Table 4). Among these sites where significant changes in amino acid characteristics occurred, the equilibrium constant (ionization of COOH) was the amino acid characteristic most affected by natural selection, which was detectable in 11 genes except for NAD6, followed by the hydrophobicity and molecular volume, which both could be detected in 10 genes.

**Table 4.**
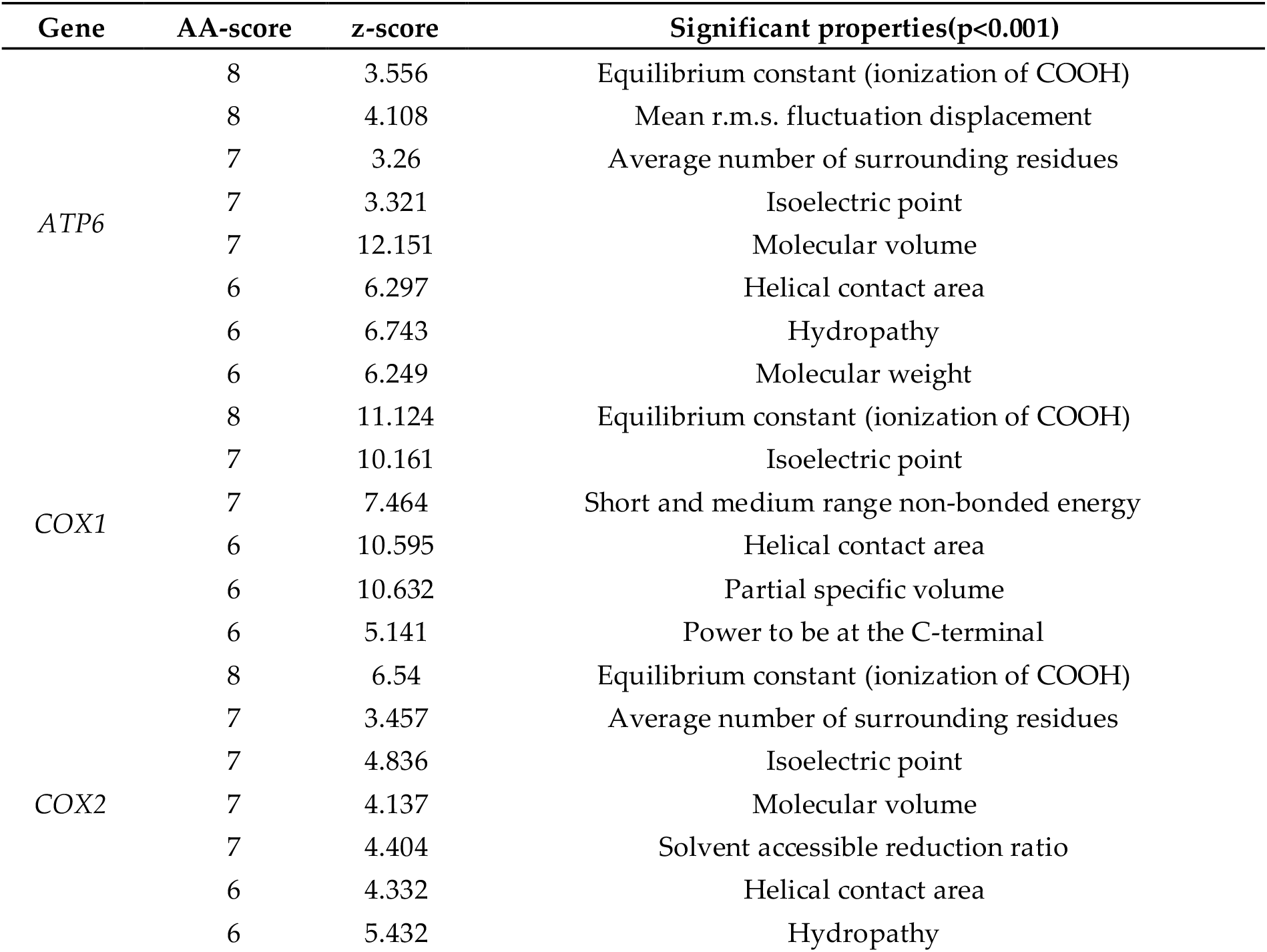
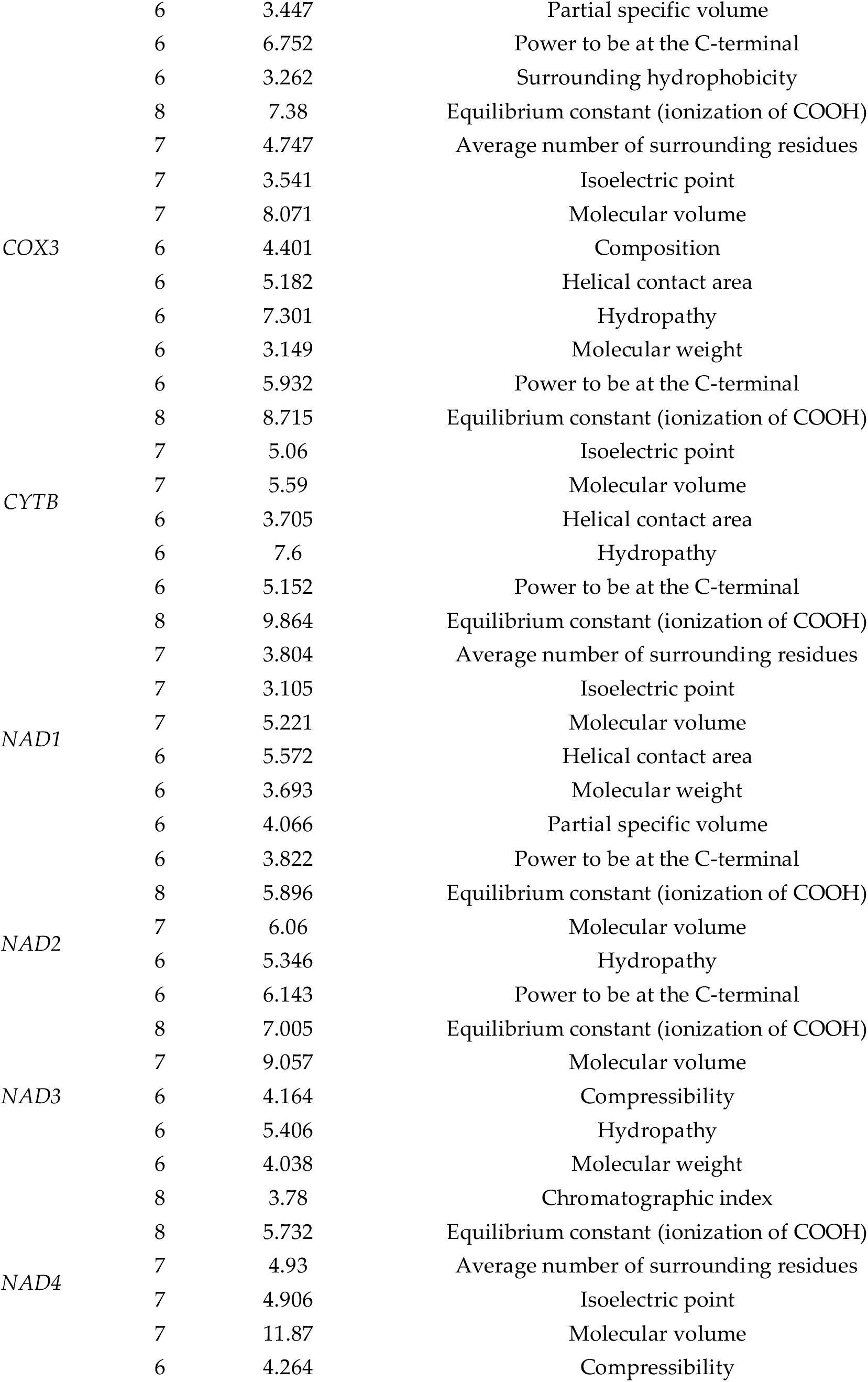
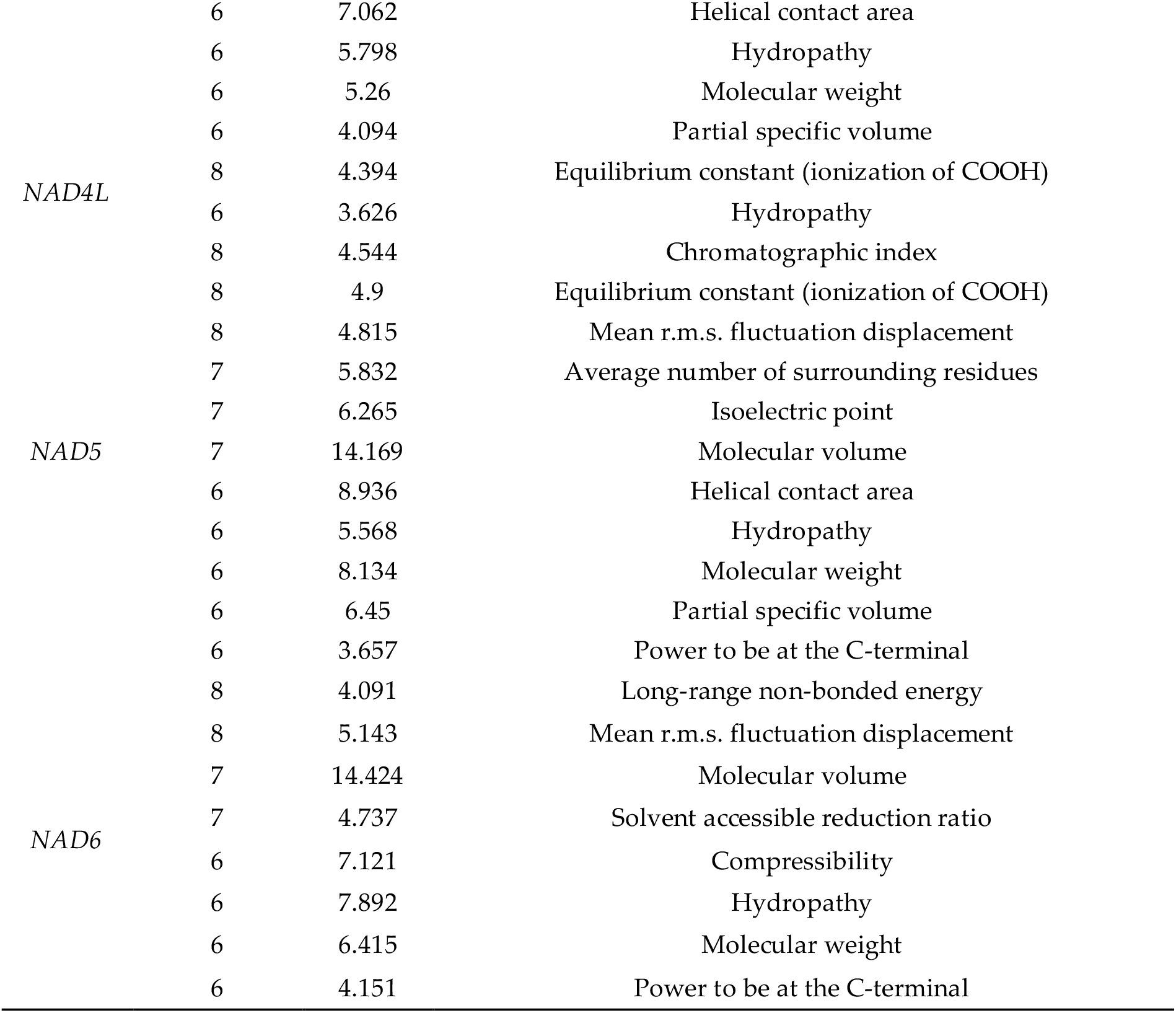
Significant changes in positive selective amino acid properties detected in TreeSAAP.

### 3.5. Phylogenetic independent contrasts analysis

To investigate the relationship between the different habitats of the turbellarians and the selection pressure of the mitochondrial PCGs, we used the Phylogenetic Independent Contrast (PIC) analysis in the ape package in R. The PIC analysis found a significant positive correlation (R = 0.61, p = 0.0033) between the ω values of the mitochondrial PCGs in turbellarians and their habitats (Fig 4). We found that different habitats significantly influenced the selection pressure of turbellarian PCGs.

**Figure 4.**
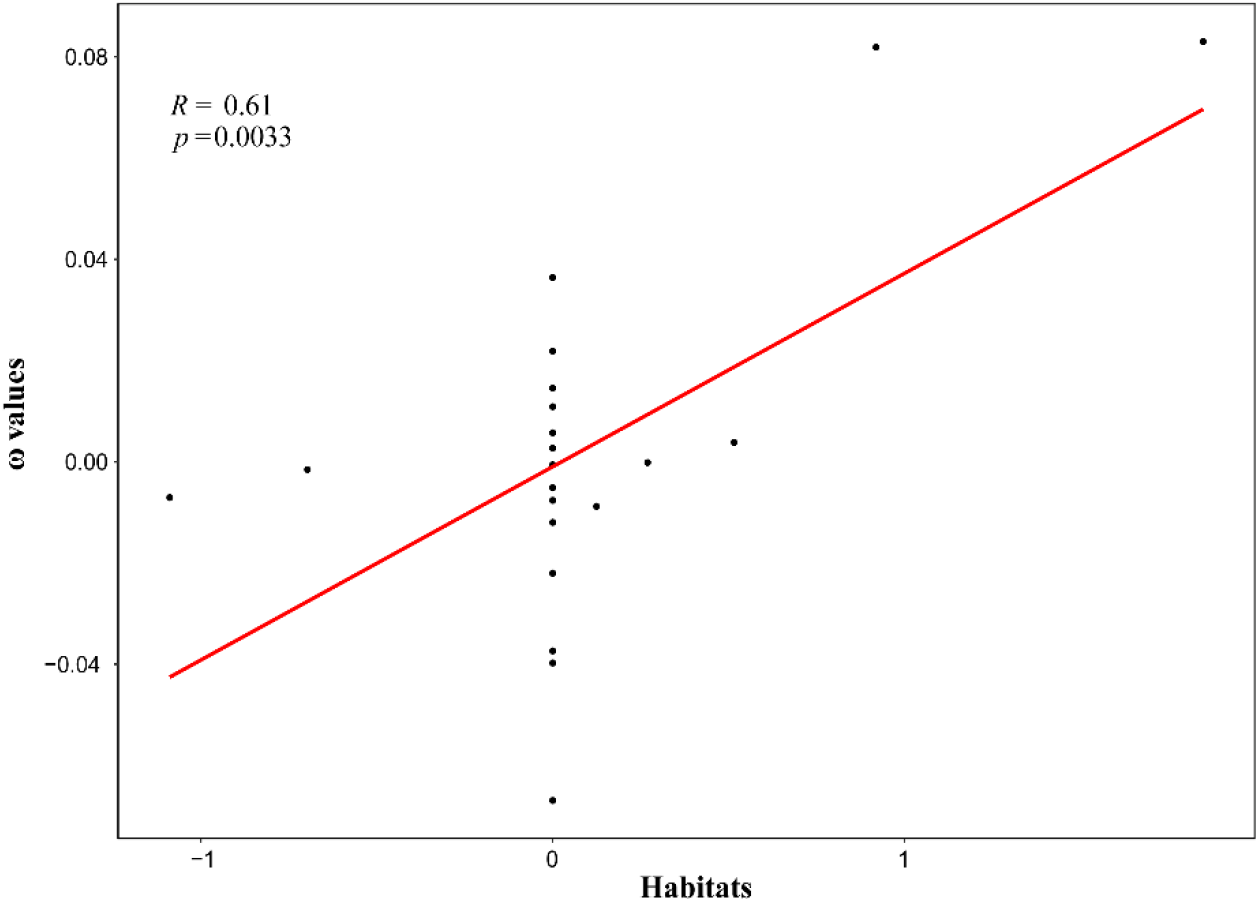
PIC analysis between habitats and ω values of protein-encoding genes in mitochondrial genomes

## 4. Discussion

The phylogenetic positions of the taxa included this study were consistent with previous inferences based on traditional taxonomy and molecular markers [17, 34]. The order Macrostomida is the earliest branch within the class Rhabditophora. Mitochondria are important organelles that provide energy for animals through oxidative phosphorylation [35]. Since animals in different ecological niches have different energy requirements, mitochondria play an important role in adapting to different ecological niches. [36].Currently, many studies focus on how animals adapt to different environments and ecological niches through the mitochondrial genome, such as cichlids[37], crickets[38], and mountain bats[39]. However, research on how turbellarians adapt to different ecological niches is very limited. Therefore, this study focuses on the adaptability of turbellarians in different ecological niches. Branch model analysis results showed that the ω values of terrestrial and aquatic turbellarians were much less than 1, which was also confirmed in the free-ratio model, indicating that purifying selection was the main evolutionary mode of mitochondrial genome in turbellarians. The result was consistent with previous research of Mustelidae[40], Tibetan loaches[37], Acrossocheilus[2], and other vertebrates[41]. Purification selection can remove harmful substitutions to maintain the normal function of PCGs[41]. Two ratio model analysis found that the ω value of three genes (ATP6, NAD2, and NAD4L) in terrestrial turbellarians were significantly higher than those in aquatic turbellarians (p <0.05), and only the COX1 was significantly lower than that in aquatic turbellarians (p <0.05), indicating that aquatic turbellarians were under more selection pressure. Comparison of ω value between different habitats using Wilcoxon test. The NAD4 in terrestrial turbellarians was significantly higher than that in aquatic turbellarians (p <0.05). The NAD4 encodes the NADH dehydrogenase of the mitochondrial gene, its product was believed to belong to the minimum assembly of core proteins that catalyze NADH dehydrogenation and electron transfer to ubiquinone (coenzyme Q10) [42] and closely related to the catalytic synthesis of ATP[12]. This indicated that the NAD4 may play an important role in the adaptation of turbellarians to terrestrial habitats.

Although purifying selection is common in mitochondrial genomes, the possibility of positive selection for individual codons cannot be ruled out, and it may promote physiological adaptation of organisms to new environments [24, 43]. In this study, codon-based selection analysis indicated that several sites in COX1, CYTB, and NAD4 of the turbellarians mitochondrial genome may be under positive selection in multiple models, but only the 77 R site of NAD4 was validated by chi-square test in the CodeML site model analysis (p < 0.05). However, only NAD4 was detected in all site model analyses, and four codons were inferred as positively selected sites for this gene. The TreeSAAP analysis of NAD4 detected 10 significant amino acid properties that underwent positive selection changes, including equilibrium constants (COOH ionization), chromatographic index, mean number of surrounding residues, isoelectric point, and hydrophobicity. Among them, the equilibrium constants (COOH ionization) property had the highest degree of change (AA-score = 8), and was detected in all 11 genes of NAD4 in the mitochondrial genome of the turbellarians. This property is an important influencing factor in protein efficiency, ROS production, and individual lifespan[44]. The study by Romero et al.[45] shows that altering the equilibrium constant can help organisms better cope with abiotic stress conditions, which are often associated with metabolic activity and ROS production[45, 46]. Changes in equilibrium constant directly affect the distribution and ability of biological species to successfully occupy new ecological niches, as well as the ability of animals to invade new ranges[45, 47]. George et al.[48] found that significant changes in the equilibrium constant (COOH ionization) affect similar regions of the genomes of amphibians, lungfish, and coelacanths, which are believed to be adaptations to increased oxygen levels and changing metabolic demands. In addition, NAD4 can participate in regulating electron transfer in the respiratory chain, providing animals with the necessary energy for activity[41, 49]. Most aquatic turbellarians have strong regenerative abilities, which may be related to the adaptive evolution of NAD4. PIC analysis shows a significant positive correlation between ω values and living environments. This indicated that different habitats significantly influenced the selection pressure of turbellarian PCGs. These results suggest that the evolution of mitochondrial-encoded protein genes is the basis for turbellarians to adapt to different living environments.

## 5. Conclusions

In summary, we investigated the evolution of mtDNA PCGs in turbellarians in different habitats. Through Wilcoxon tests, selection pressure, and PIC analysis, purifying selection was found to be the main evolutionary pattern of turbellarian mtDNA genomes. The genes ATP6, NAD2, and NAD4L were identified experienced higher selection pressures in aquatic turbellarians, while COX1 was experienced higher selection pressures in terrestrial turbellarians. Furthermore, positive selection sites were detected in NAD4 in all site models. Additionally, equilibrium constant (ionization of COOH) was the amino acid characteristic with the highest degree of variation in NAD4, indicating its important role in the adaptive evolution of turbellarians to different habitats. The adaptive evolution of mtDNA-encoded protein genes serves as the basis for turbellarians to adapt to different habitats.

## Supplementary Materials

Table Supplementary

## Author Contributions

R.L. wrote the manuscript. P.C designed the study. R.L. and H.F. carried out the experiment. J.W. and P.C. provided resources and technical support. All authors reviewed the manuscript. All authors have read and agreed to the published version of the manuscript.

## Data Availability

All genomic data used in this study were downloaded from publicly available repository NCBI (https://www.ncbi.nlm.nih.gov/) as detailed in the Supplementary material.

