## Supplementary material for "Natural selection and evolution of mitochondrial genome in terrestrial and aquatic turbellarians": Table Supplementary

**Table S1.** List of species used in this study

| Order | Family | Species | Accession number | Length |
|---|---|---|---|---|
| Macrostomida | Macrostomidae | Macrostomum lignano | NC_035255 | 14193 |
| Polycladida | Hoploplanidae | Hoploplana elisabelloi | NC_028200 | 15206 |
| | Planoceridae | Planocera reticulata | NC_036051 | 14724 |
| | Prosthiostomidae | Enchiridium sp. MTA_2015 | NC_028199 | 14651 |
| | | Prosthiostomum siphunculus | NC_028201 | 15181 |
| Rhabdocoela | Umagillidae | Syndesmis echinorum | NC_050392 | 15053 |
| | | Syndesmis kurakaikina | MT063057 | 14226 |
| Tricladida | Dugesiidae | Girardia tigrina | MW972220 | 15938 |
| | | Girardia sp. ER-2015 | KP090061 | 15951 |
| | | Dugesia japonica | AB618487 | 17799 |
| | | Schmidtea mediterranea | NC_022448 | 27133 |
| | Geoplanidae | Amaga expatria | NC_057980 | 14962 |
| | | Bipalium adventitium | MZ561467 | 15494 |
| | | Bipalium kewense | NC_045216 | 15666 |
| | | Bipalium vagum | MZ561468 | 17149 |
| | | Diversibipalium multilineatum | MZ561469 | 15660 |
| | | Diversibipalium sp. MNHN JL281 | MZ561470 | 15989 |
| | | Humbertium sp. MNHN JL090 | MZ561471 | 15524 |
| | | Humbertium sp. MNHN JL351 | MZ561472 | 15540 |
| | | Obama sp. MAP-2014 | KP208777 | 14909 |
| | | Parakontikia ventrolineata | MT081960 | 17210 |
| | | Platydemus manokwari | MT081580 | 19959 |
| | Uteriporidae | Miroplana shenzhensis | NC_062124 | 14344 |
| | | Obrimoposthia wandeli | NC_050050 | 15185 |
| Rhinebothriidea | Rhinebothriidae | Rhinebothrium reydai | NC_044703 | 13506 |
| Gyrodactylidea | Gyrodactylidae | Gyrodactylus salaris | NC_008815 | 14790 |
| Strigeidida | Schistosomatidae | Schistosoma mekongi | NC_002529 | 14072 |

**Table S2.** Analysis of 12 protein-coding genes of turbellarian by CodeML program branching model

| Gene | Model | Lnl | np | ω | Test | LRT | df | p-value |
|---|---|---|---|---|---|---|---|---|
| *ATP6* | One Model | -11742.98 | 47 | 0.03349 | - | - | - | - |
| | Free Model | -11676.79 | 91 | various | Free x One | 132.38 | 44 | 0 |
| | Two model | -11740.7 | 48 | ω0: 0.04155<br>ω1: 0.02375 | Two x One | 4.57 | 1 | 0.033* |
| *COX1* | One Model | -22727.12 | 47 | 0.02824 | - | - | - | - |
| | Free Model | -22538.88 | 91 | various | Free x One | 376.48 | 44 | 0 |
| | Two Model | -22723.33 | 48 | ω0: 0.02463<br>ω1: 0.03408 | Two x One | 7.58 | 1 | 0.006** |
| *COX2* | One Model | -14936.78 | 47 | 0.07005 | - | - | - | - |
| | Free Model | -14846.76 | 91 | various | Free x One | 180.04 | 44 | 0 |
| | Two Model | -14936.75 | 48 | ω0: 0.06905<br>ω1: 0.07205 | Two x One | 0.06 | 1 | 0.81 |
| *COX3* | One Model | -13369.47 | 47 | 0.03307 | - | - | - | - |
| | Free Model | -13267 | 91 | various | Free x One | 204.93 | 44 | 0 |
| | Two Model | -13368.65 | 48 | ω0: 0.03073<br>ω1: 0.0416 | Two x One | 1.65 | 1 | 0.2 |
| *NAD1* | One Model | -13518.05 | 47 | 0.04036 | - | - | - | - |
| | Free Model | -13435.07 | 91 | various | Free x One | 165.95 | 44 | 0 |
| | Two Model | -13517.83 | 48 | ω0: 0.03883<br>ω1: 0.04296 | Two x One | 0.43 | 1 | 0.512 |
| *NAD2* | One Model | -16967.73 | 47 | 0.04984 | - | - | - | - |
| | Free Model | -16889.49 | 91 | various | Free x One | 156.48 | 44 | 0 |
| | Two Model | -16964.61 | 48 | ω0: 0.06412<br>ω1: 0.02258 | Two x One | 6.24 | 1 | 0.012* |
| *NAD3* | One Model | -5488.746 | 47 | 0.06551 | - | - | - | - |
| | Free Model | -5458.705 | 91 | various | Free x One | 60.08 | 44 | 0.054 |
| | Two Model | -5488.733 | 48 | ω0: 0.06458<br>ω1: 0.06684 | Two x One | 0.03 | 1 | 0.873 |
| *NAD4* | One Model | -23027.05 | 47 | 0.04288 | - | - | - | - |
| | Free Model | -22942.48 | 91 | various | Free x One | 169.15 | 44 | 0 |
| | Two Model | -23026.87 | 48 | ω0: 0.0444<br>ω1: 0.0402 | Two x One | 0.37 | 1 | 0.542 |
| *NAD4L* | One Model | -4937.277 | 47 | 0.03428 | - | - | - | - |
| | Free Model | -4903.01 | 91 | various | Free x One | 68.53 | 44 | 0.010* |
| | Two Model | -4931.32 | 48 | ω0: 0.07146<br>ω1: 0.01658 | Two x One | 11.91 | 1 | 0.00056*** |
| *NAD5* | One Model | -27704.7 | 47 | 0.04921 | - | - | - | - |
| | Free Model | -27567.8 | 91 | various | Free x One | 273.81 | 44 | 0 |
| | Two Model | -27704.09 | 48 | ω0: 0.05226<br>ω1: 0.04487 | Two x One | 1.22 | 1 | 0.27 |
| *NAD6* | One Model | -8895.456 | 47 | 0.0503 | - | - | - | - |
| | Free Model | -8827.486 | 91 | various | Free x One | 135.94 | 44 | 0 |

| | | | | | | | | |
|---|---|---|---|---|---|---|---|---|
| | Two Model | -8894.003 | 48 | ω0: 0.06086<br>ω1: 0.03642 | Two x One | 2.9 | 1 | 0.088 |
| *CYTB* | One Model | -17544.49 | 47 | 0.03438 | - | - | - | - |
| | Free Model | -17430.1 | 91 | various | Free x One | 228.79 | 44 | 0 |
| | Two Model | -17544.44 | 48 | ω0: 0.03374<br>ω1: 0.03555 | Two x One | 0.11 | 1 | 0.737 |

**Table S3.** Codon positions under positive selection detected by branch-site model using codeML

| Gene | Model | Lnl | Test | p-value | Positive site |
|---|---|---|---|---|---|
| *ATP6* | Model A | -7671.182954 | Model A x Model A null | 1 | 20 I 1.000**,27 Q 1.000**,35 T 0.974*,41 I 0.988*, 45 R 0.998**, 52 S 0.998**, 66 S 0.960*, 94 S 0.987*, 118 F 0.981*, 136 I 0.976*, 137 V 1.000**, 141 E 1.000** |
| | Model A null | -7671.182954 | | | |
| *COX1* | Model A | -19498.9504 | Model A x Model A null | 1 | 21 S 0.998**, 31 S 0.993**, 73 G 0.983*, 74 I 1.000**, 89 L 0.980*, 92 I 0.979*, 117 S 1.000**, 119 V 0.999**, 149 M 1.000**, 150 K 0.971*, 238 F 1.000**, 303 D 1.000**, 304 N 0.994**, 305 S 0.953*, 307 S 0.990**, 311 L 0.998**, 370 V 0.979*, 371 R 0.993**, 374 P 1.000**, 375 F 0.999**, 378 F 0.967*, 407 V 1.000**, 412 E 0.991**, 417 K 1.000**, 440 E 0.992**, 441 E 0.998**, 449 S 1.000**, 454 V 0.956*, 456 I 0.999**, 461 G 0.975*, 466 Y 1.000** |
| | Model A null | -19498.9504 | | | |
| *COX2* | Model A | -9349.341978 | Model A x Model A null | 1 | 17 C 0.997**, 26 I 0.994**, 29 N 0.997**, 33 N 0.999**, 54 F 0.999**, 68 S 0.997**, 70 V 1.000**, 71 L 1.000**, 86 F 0.983*, 113 K 0.968*, 115 L 0.980*, 145 Q 0.957*, 147 F 0.989*, 153 S 0.951*, 176 F 0.998** |
| | Model A null | -9349.341978 | | | |
| *COX3* | Model A | -10603.41775 | Model A x Model A null | 1 | 14 I 0.997**, 23 I 0.990**, 25 V 0.986*, 28 F 0.973*, 36 S 1.000**, 43 V 0.988*, 51 V 0.995**, 127 S 0.980*, 131 L 1.000**, 135 S 0.985*, 178 H 0.952* |
| | Model A null | -10603.41775 | | | |
| *NAD1* | Model A | -9984.778988 | Model A x Model A null | 1 | 136 H 0.961*, 137 G 0.997** |
| | Model A null | -9984.778988 | | | |
| *NAD2* | Model A | -9794.139459 | Model A x Model A null | 0.998 | 18 F 0.971*, 20 E 1.000**, 42 E 0.995**, 62 K 0.998**, 69 F 0.978*, 133 C 0.973* |
| | Model A null | -9794.139461 | | | |
| *NAD3* | Model A | -3952.104473 | Model A x Model A null | 1 | 61 F 0.975* |
| | Model A null | -3952.104473 | | | |

| | | | | | |
|---|---|---|---|---|---|
| *NAD4* | Model A<br>Model A null | -15191.08918<br>-15191.08918 | Model A x Model A null | 1 | 73 R 0.989*, 244 S 0.997**, 250 S 0.987* |
| *NAD4L* | Model A<br>Model A null | -3253.434694<br>-3253.434694 | Model A x Model A null | 1 | NA |
| *NAD5* | Model A<br>Model A null | -16634.17391<br>-16634.17391 | Model A x Model A null | 1 | 15 R 0.999**, 78 S 0.987*, 103 Q 0.985*, 161 R 1.000**, 162 F 0.999**, 205 S 0.968* |
| *NAD6* | Model A<br>Model A null | -3046.357803<br>-3046.357803 | Model A x Model A null | 1 | NA |
| *CYTB* | Model A<br>Model A null | -14593.57919<br>-14593.57919 | Model A x Model A null | 1 | 1 S 0.997**, 192 L 0.996**, 200 L 0.959*, 210 L 0.979*, 277 F 0.997**, 298 R 0.975*, 306 L 0.981* |
